# Traveling waves in attention networks inherit the rhythms of sensory systems

**DOI:** 10.64898/2026.11.04.763906

**Authors:** Yue Kong, Kirsten Petras, David M. Alexander, Benjamin Rohaut, Laura Dugué

## Abstract

Attention emerges from coordinated activity across distributed brain regions^1–6^, each with their own intrinsic rhythm: alpha (7-12Hz) in visual cortex^7–12^ and theta (3-7Hz) in auditory cortex^13–15^. When attention is sustained on a visual location, alpha power increases over the corresponding ipsilateral hemisphere, a robust neurophysiological marker of sustained visuospatial attention named alpha lateralization^16–20^. Hemispheric asymmetry during sustained attention was recently also reported for anterior-to-posterior alpha traveling waves^21,22^ –smooth phase shifts across the cortex^23,24^. This finding suggests that traveling waves support top-down inter-areal communication within the attention network. Here we asked whether traveling waves indeed reflect top-down attention signals or are merely a byproduct of sustained visuospatial attention. We manipulated attention through different sensory modalities (auditory cueing and visual stimulation), thus engaging different sensory cortices, while maintaining the task as visual discrimination. Using electroencephalography, we tested the hypothesis that if traveling waves reflect attention signals within the network, their frequency should follow the intrinsic rhythm of the engaged sensory system: theta for auditory cueing and alpha for visual stimulation. If instead they are a byproduct of attention allocated to visual information, their frequency should remain in alpha. Our results support the former: auditory cueing led to lateralized theta traveling waves, while visual stimulation produced lateralized alpha traveling waves. Crucially, traveling-wave lateralization in both frequencies correlated with trial-by-trial alpha power lateralization. Together, attention signals appear to propagate as waves at the intrinsic rhythm of the engaged sensory system and locally modulate alpha power lateralization in task-relevant visual regions.

## Results

Sustained spatial attention to a visual stimulus is correlated with a well-characterized brain signature: alpha power, the intrinsic rhythm of visual cortex, increases over the hemisphere ipsilateral to the attended location^16–20^. Recent studies have shown that alpha traveling waves propagating from anterior to posterior brain regions are similarly lateralized^21,22^. This observation suggests that traveling waves support top-down inter-areal communication within the attention network (see^25^), which we test here using electroencephalography (EEG).

Human participants (N=18) performed an orientation discrimination task in which an auditory cue directed visual attention towards the upcoming visual target (grating patch) location (**Figure 1A**; cue validity: 66.6%, see STAR Methods). Perceptual sensitivity (d’) was significantly higher in valid than invalid trials (mean ± Standard Error of the Mean (SEM): 2.044 ± 0.169 and 0.479 ± 0.119; one-sided paired t-test, *t*(17) = 6.907, *p* < 0.001, Cohen’s *d* = 1.628; **Figure 1B**), whereas median reaction time did not differ between conditions (two-sided paired t-test, *t*(17) = -0.709, *p* = 0.488, Cohen’s *d* = -0.167), excluding a speed-accuracy trade-off^26,27^. These behavioral results confirm that auditory cueing successfully manipulated visuospatial attention and that attentional orienting improved perceptual sensitivity.

**Figure 1.**
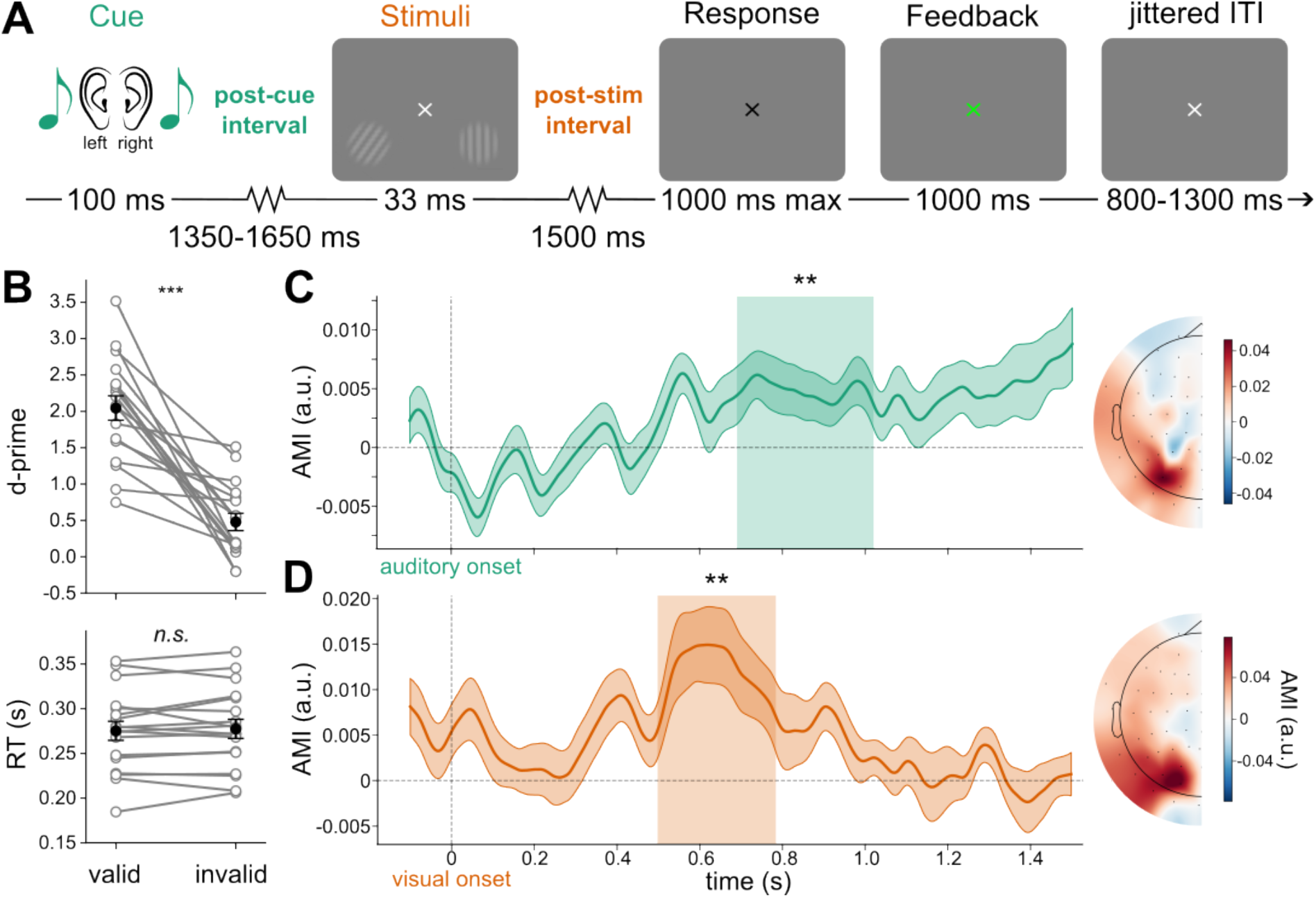
Auditory cues and visual stimuli directed visuospatial attention and induced alpha power lateralization. **A** Attentional cueing paradigm. A transient, spatially informative, binaural auditory cue (a sine- or semisine-formed pure tone) instructed participants to orient their attention to either the left or right lower quadrant. After a jittered delay, two grating patches were presented. After a delay of 1500 ms, participants reported the orientation of the target grating (tilted; the vertical grating is the distractor), followed by feedback (red/green). Valid trials (66.6%): target at cued location (orienting). Invalid trials (33.3%): target at uncued location (reorienting). Eye-tracking was used throughout to ensure central fixation and avoid signal artifacts. ITI: inter-trial interval. **B** Perceptual sensitivity (d’, top) and median reaction time (RT, bottom) in valid and invalid trials. Gray: individual participants; black: group mean ± SEM. ***: *p* < 0.001, one-sided paired t-test; *n.s.*: non-significant, two-sided paired t-test. **C** Time-resolved AMI in the post-auditory cue period. Solid line: group mean; shaded band: SEM. The shaded rectangle marks the significant cluster (0.690-1.020 s, ** *p* = 0.008, one-tailed cluster-based permutation test). Right panel: topography of AMI averaged within the significant cluster; positive values indicate higher alpha power in ipsilateral hemisphere. **D** same as **C**, but in the post-visual stimulus period (significant cluster: 0.498-0.784 s; ** *p* = 0.006).

In the remainder of the analysis, we focused on sustained attentional orienting within two periods of interest: (1) post-auditory cue period in which sustained visual attention was directed by an auditory cue, and (2) post-visual stimulus period in which sustained attention was directed by a visual stimulus^19,20^. Only valid trials were included in the analysis of post-visual stimulus period.

### Alpha power lateralized for both auditory and visual attentional manipulation

To assess alpha power lateralization, we computed a time-resolved Attention Modulation Index (AMI; per trial, frequency and time point) as the normalized difference between ipsilateral and contralateral alpha power (7-12Hz; using all electrodes except for midline; see STAR Methods). The AMI was significantly greater than zero during both the post-auditory cue period (cluster-based permutation test: 0.690-1.020 s, *p* = 0.008; **Figure 1C**, left panel) and the post-visual stimulus period (0.498-0.784 s, *p* = 0.006; **Figure 1D**, left panel). Topographies of AMI within these significant time windows show that alpha lateralization was most pronounced over occipital electrodes (**Figure 1C and 1D**, right panels). Together these results confirm that alpha power lateralizes when attention is sustained to a visual location, both after auditory and visual attentional manipulation.

### Sensory events induced traveling waves at distinct temporal frequencies

To characterize traveling waves on single trials, we compared the spatial patterns of phase dynamics with data-driven templates of wave propagations (**Figure 2A**; see STAR Methods). When a wave propagates across the scalp, its phase shifts smoothly and systematically over space. Singular value decomposition (SVD), a data-driven method that decomposes the phase data into spatial components, revealed that the dominant components in our data resembled one-cycle rotating waves in the sagittal plane (**Figure S1**), in line with previous reports of attention-related traveling waves propagating predominantly from anterior to posterior on the dorsal scalp^21,22,25^. To facilitate interpretation and comparison across participants and conditions^25^, we constructed rotating wave templates, in which phase propagates uniformly around a central point (i.e., the singularity) for a full cycle. These templates provided a common reference, identical across participants and conditions and independent of any subset of data, consistent with the primary component from SVD. They also defined two propagation directions as exact opposites (complex conjugates of each other): clockwise (CW) and counterclockwise (CCW) rotation, both viewed from the left (**Figure 2A**). Because the templates were defined in the same sagittal plane for all electrodes, CW denotes anterior-to-posterior and CCW posterior-to-anterior propagation over the dorsal scalp in both hemispheres. Wave activity in each direction was quantified on each trial and at each time point as the similarity between the measured phase pattern and the corresponding template (ranging from 0 to 1, where 1 indicates a perfect match). We then used the difference between the two directions (CW – CCW) to quantify the net direction of propagation: positive values indicate a predominance of anterior-to-posterior waves and negative values a predominance of posterior-to-anterior waves.

**Figure 2.**
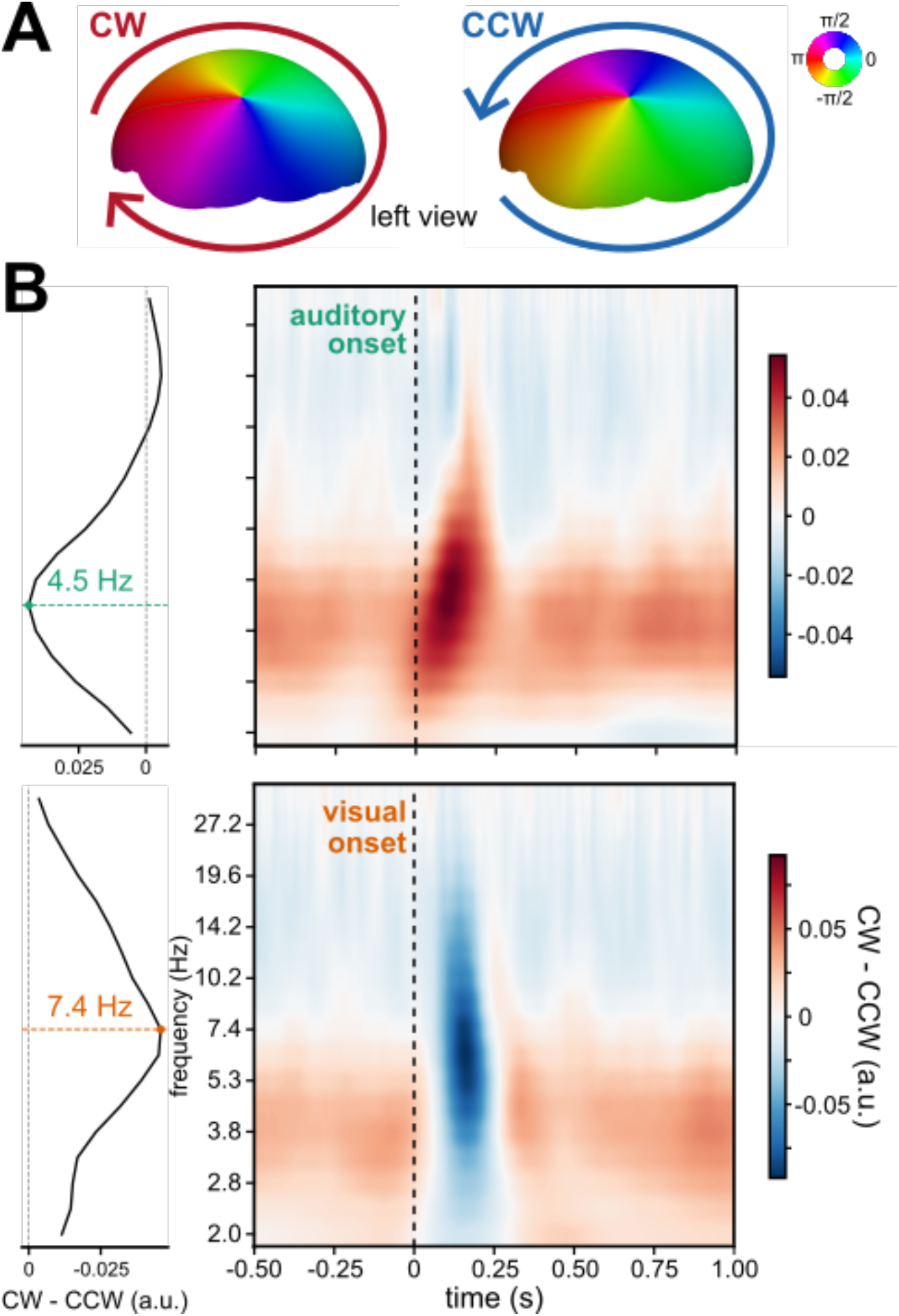
Auditory and visual onsets induced traveling waves at distinct temporal frequencies. **A** Rotational templates used to quantify traveling wave activity, shown in the left view. Color indicates oscillatory phase. Phase propagates uniformly around a central singularity in the clockwise (CW, left) or counterclockwise (CCW, right) direction, completing one full cycle per rotation. Over the dorsal scalp, CW and CCW rotation correspond to anterior-to-posterior and posterior-to-anterior propagation, respectively. **B** Time-frequency representation of directional wave activity (CW - CCW) after auditory (top) and visual onsets (bottom). Red, CW waves dominate; blue, CCW waves dominate. Left and right panels share the same frequency axis (log-scaled). Left, frequency profiles averaged over 0-0.2 s after onset; dashed lines mark the peak frequencies (auditory: 4.5 Hz; visual: 7.4 Hz).

Auditory cues induced a transient increase in CW (anterior-to-posterior) wave activity centered in the theta frequency band (peak at 4.5 Hz, averaged over 0-0.2 s after cue onset), whereas the visual target induced a transient increase of CCW (posterior-to-anterior) wave activity centered in the alpha band (peak at 7.4 Hz, averaged over 0-0.2 s after stimulus onset; **Figure 2B**). Note that low-theta CW traveling waves were present throughout the trial, but here we only focus on the stimulus-induced waves.

### CW Traveling waves lateralized at modality-specific frequencies

Having established that auditory cueing and visual stimulation evoked traveling waves at distinct temporal frequencies, we next asked whether traveling waves were modulated by visuospatial attention in the same way as occipital alpha power. We therefore focused on the temporal frequency band of the wave induced by each sensory event: theta (3-7 Hz) for the post-auditory cue period and alpha (7-12 Hz) for the post-visual stimulus period (**Figure 2B**). To assess the hemispheric asymmetry of traveling waves, we restricted the whole-brain rotational templates to the electrodes of each hemisphere and fitted them separately to the data of that hemisphere (see STAR Methods). Because the templates rotate around the interaural axis, each hemisphere’s template retained the full rotation cycle, and the shapes of the left and right templates were identical. CW and CCW wave activity was then averaged within the time windows showing significant alpha power lateralization (0.690-1.020 s after auditory cue onset; 0.498-0.784 s after visual stimulus onset; **Figure 1**).

For the post-auditory cue period, a 2×2 repeated-measures analysis of variance (ANOVA) on theta-band wave activity, with factors hemisphere (ipsilateral vs. contralateral to the attended location) and direction (CW vs. CCW), revealed a significant interaction (*F*(1,17) = 5.882, *p* = 0.027; **Figure 3A**, left panel). This interaction was primarily driven by more CW waves (i.e., anterior-to-posterior) in the ipsilateral than contralateral hemisphere (*t*(17) = 2.676, *p_Bonferroni* = 0.032, Cohen’s *d* = 0.631); the hemispheric difference for CCW waves was not significant (*t*(17) = -1.933, *p_Bonferroni* = 0.140, Cohen’s *d* = -0.456). A significant main effect of direction (*F*(1,17) = 26.206, *p* < 0.001) indicated that across hemispheres, CW waves were more prevalent than CCW waves in the theta band. In contrast, across directions, theta-band wave activity did not differ between the ipsilateral and contralateral hemispheres (*F*(1,17) = 0.710, *p* = 0.411).

**Figure 3.**
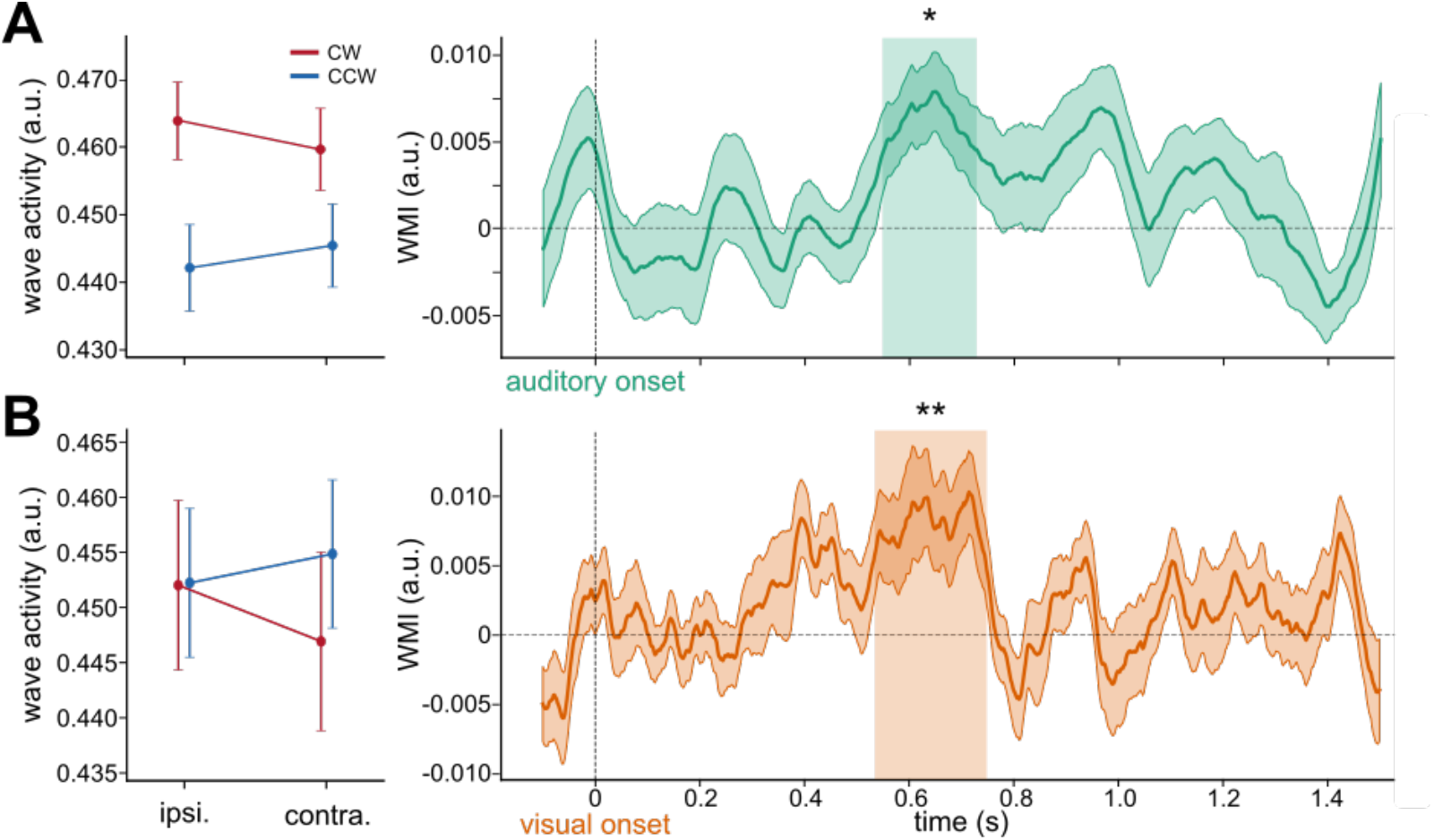
CW traveling wave lateralized at modality-specific frequencies. **A** Theta-band CW waves lateralized in the post-auditory cue period. Left panel, CW (red) and CCW (blue) wave activity in ipsilateral and contralateral hemispheres, averaged over the significant time window of alpha power lateralization (0.690-1.020 s; see Figure 1C). Error bars, between-participant SEM. A 2×2 repeated-measures ANOVA revealed a significant hemisphere x direction interaction (*F*(1,17) = 5.882, *p* = 0.027); CW waves were stronger in the ipsilateral hemisphere, whereas CCW waves did not lateralize. Right panel, time-resolved WMI of CW theta waves (time 0, cue onset). Solid line: group mean; shaded band: SEM. The shaded rectangle marks the significant cluster (0.548-0.728 s, * *p* = 0.019, one-tailed cluster-based permutation test). **B** same as **A**, but for alpha-band traveling waves in the post-visual stimulus period. Left panel, wave activity averaged over 0.498-0.784 s (see Figure 1D); hemisphere x direction interaction, *F*(1,17) = 12.238, *p* = 0.003. Right panel, LI of CW alpha waves; significant cluster, 0.534-0.748 s, ** *p* = 0.003.

For the post-visual stimulus period, the same ANOVA on alpha-band wave activity revealed a significant interaction effect (*F*(1,17) = 12.238, *p* = 0.003; **Figure 3B**, left panel), with neither main effect reaching significance (direction: *F*(1,17) = 0.319, *p* = 0.580; hemisphere: *F*(1,17) = 2.504, *p* = 0.132). Again, the interaction was driven primarily by more CW waves in the ipsilateral hemisphere (*t*(17) = 3.388, *p_Bonferroni* = 0.007, Cohen’s *d* = 0.798), whereas the hemispheric difference for CCW waves was not significant (*t*(17) = -2.225, *p_Bonferroni* = 0.080, Cohen’s *d* = -0.524).

In both post-auditory cue and post-visual stimulus periods, attention thus increased CW waves in the hemisphere ipsilateral to the attended location, paralleling the ipsilateral increase in alpha power and consistent with previous reports of lateralized anterior-to-posterior alpha waves during sustained attention^21^. We therefore focused subsequent analyses on CW waves.

Because the analyses above were restricted to the time windows of alpha power lateralization, we next examined when CW wave lateralization appeared, independently of these windows. To this end, we computed a Wave Modulation Index (WMI) analogous to the AMI (see STAR Methods). The WMI of CW theta waves was significantly greater than zero 0.548-0.728 s after auditory onset (cluster-based permutation test, *p* = 0.019; **Figure 3A**, right panel), and the WMI of CW alpha waves was significantly greater than zero 0.534-0.748 s after visual onset (*p* = 0.003; **Figure 3B**, right panel). Thus, in both periods, CW (anterior-to-posterior) waves were lateralized in the same manner as alpha power, but at a temporal frequency that matched the waves induced by the attention-directing sensory event.

### Traveling wave lateralization correlated with alpha power lateralization

So far, we identified a dissociation between two attentional signatures: 1) alpha power lateralization in both post-auditory cue and post-visual stimulus periods; 2) attentional modulation (lateralization) of CW waves at distinct frequencies: theta for auditory manipulation and alpha for visual manipulation. We last asked whether traveling-wave lateralization was correlated with alpha power lateralization at the single-trial level. The two measures capture distinct properties of the same signals: the WMI was computed from phase alone, after normalizing the wavelet coefficients to unit magnitude, and thus reflects the spatial organization of phase across electrodes, whereas the AMI was computed from oscillatory power and thus reflects amplitude only.

For each participant, we correlated WMI with AMI across trials, both averaged over the union of the significant wave and power clusters (0.548-1.020 s after auditory cue onset; 0.498-0.784 s after visual stimulus onset). We tested the Fisher z-transformed correlation coefficients against zero. For the post-auditory cue period, CW theta wave lateralization correlated positively with alpha power lateralization; although modest, this correlation was consistent across participants (**Figure 4A**; mean ± SD *r* = 0.039 ± 0.065, 13/18 participants with *r* > 0; two-sided one-sample t test, *t*(17) = 2.532, *p* = 0.022, Cohen’s *d* = 0.597). For the post-visual stimulus period, CW alpha wave lateralization likewise correlated with alpha power lateralization (**Figure 4B**; mean ± SD *r* = 0.198 ± 0.111, 17/18 participants with *r* > 0; *t*(17) = 7.373, *p* < 0.001, Cohen’s *d* = 1.738). The stronger correlation after visual stimulation may partly reflect the shared frequency band of alpha, because phase estimates are more reliable when oscillatory amplitude is high. The cross-frequency correlation after auditory cues, although weaker, is less susceptible to this dependence.

**Figure 4.**
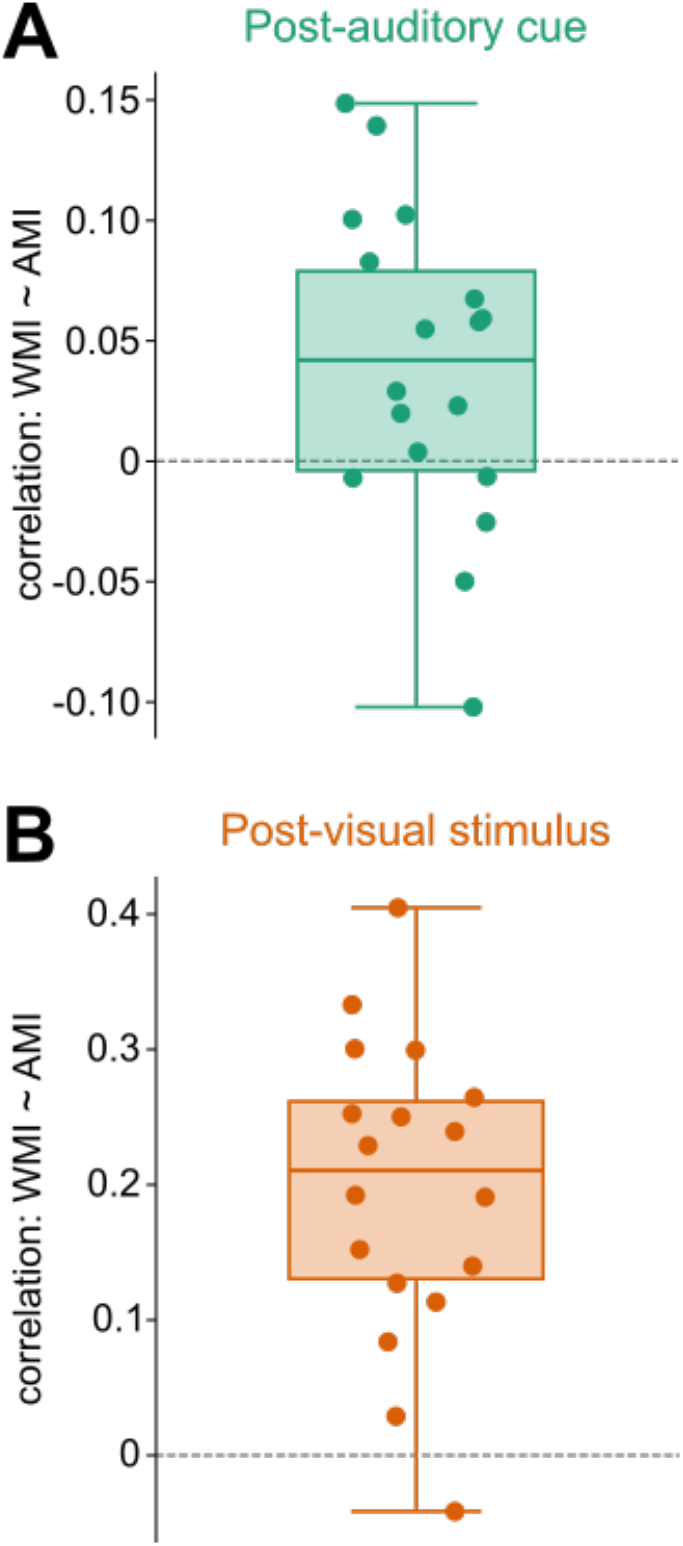
Traveling wave lateralization correlated with alpha power lateralization on a trial-by-trial basis. **A** Within-participant correlations between the WMI of CW theta waves and the AMI in the post-auditory cue period, both averaged over 0.548-1.020 s. Each dot represents the Pearson’s *r* of one participant (total N = 18). Box plots show the median and interquartile range (IQR); whiskers extend to the most extreme values within 1.5 × IQR. Two-sided one-sample t-test on Fisher z-transformed coefficients showed correlation coefficients differed significantly from zero, *t*(17) = 2.532, *p* = 0.022. **B** same as **A**, but for CW alpha waves in the post-visual stimulus period, averaged over 0.498-0.784 s. One-sample t-test showed *t*(17) = 7.373, *p* < 0.001.

## Discussion

Our results indicate that traveling waves in attention networks inherit the rhythm of the engaged sensory system. This study investigated whether traveling waves reflect top-down attention signals by separating the sensory systems of the attentional signal and its target. We found a dissociation between two attentional signatures: traveling waves lateralized in a frequency band that depended on the sensory modality of the attentional manipulation (theta for auditory and alpha for visual), whereas alpha power lateralized consistently across modalities. Moreover, traveling-wave lateralization correlated with alpha power lateralization on a trial-by-trial basis. Together these findings suggest that traveling waves reflect top-down attention signals, specific to the sensory modality, instead of being merely a byproduct of sustained visuospatial attention.

Attention emerges from the coordinated activity of distributed brain regions. Specifically, sustained attention relies on frontoparietal regions^1–3^, which interact with sensory cortices and subcortical structures such as the thalamus^4–6^. Within this network, distinct regions have been associated with neural oscillations at distinct temporal frequencies. In occipital visual cortex, alpha oscillations naturally emerge^7,12,28,29^ and are associated with perception and attention. Auditory cortex, in contrast, exhibits endogenous theta-band rhythms^13–15^, associated with auditory perception and speech processing^30–32^. Frontal regions, when involved in visual attention tasks, generally show slower rhythms than those of visual cortex^33–35^. These region-specific frequencies likely depend on the neurophysiological properties of each region, i.e., the integration time constants and conduction delays^9–11,24,36^, consistent with the characteristic spectral profiles of cortical regions at resting state^37^ and with the systematic decrease of intrinsic frequencies along the posterior-anterior axis^11,35^.

Traveling waves have been proposed to support inter-areal communication within large-scale brain networks^23–25,38^. Sustained visual attention lateralizes anterior-to-posterior traveling waves at alpha frequency bands^21^; inducing such waves in one hemisphere improves attentional performance in the ipsilateral visual field^22^. Because previous studies relied on visual stimuli, the observed waves likely reflected communication between frontal and occipital regions of the attention network, at the intrinsic rhythm of visual cortex, alpha. Here, by directing visual attention with auditory cues, we engaged an additional sensory system in the attention network and thereby altered the path of inter-areal communication. Accordingly, traveling waves were lateralized at the theta frequency, indicating that their frequency depends on the regions engaged in the communication. Moreover, traveling wave lateralization and alpha power lateralization were correlated on a trial-by-trial basis, as would be expected if traveling waves reflected the attentional signals to visual cortex. After auditory cueing, this correlation was small, as is typical for single-trial EEG measures, but it was consistent across participants and held across frequencies, making it unlikely to arise from a shared dependence of phase and amplitude estimates within the same band. Therefore, traveling waves are not merely a byproduct of sustained visuospatial attention but reflect attention signaling within the attention network.

This principle may also account for the different temporal frequencies reported in previous literature for sustained and exploratory attention –alpha for sustained^16–18,39^ and theta for exploratory^20,40,41^. Indeed, different attentional states likely recruit different paths through the attention network: sustained visual attention was proposed to involve top-down signals from frontal to occipital regions, whereas exploratory attention would heavily rely on frontal regions to sequentially shift attention from location to location, which may explain its slower theta frequency^25,42–47^. This view aligns with the recent proposal that global traveling waves can emerge from interactions within networks while being constrained by region-specific neurophysiological properties^24^. Together, we argue that by manipulating different nodes within the attention network (visual, auditory and frontal cortices) and then observing changes in the temporal frequency of traveling waves, we can furnish evidence that the frequency of attentional traveling waves reflects the communication paths within the attention network, not a fixed property of attention itself.

A recurring concern is that traveling waves observed non-invasively may not reflect activity propagating across the cortex, but could instead arise from a few spatially discrete, phase-lagged sources whose signals are mixed at the sensors through volume conduction^48,49^. Two considerations mitigate this concern. First, a previous study using direct intracranial recordings from gray matter (not subject to volume conduction by the scalp and other intervening tissues) showed that cortical dynamics are dominated by low spatial frequencies^50^, in line with the large-scale traveling waves studied here. Second, Dugué and Chavane^24^ recently argued that apparent phase gradients on the scalp can emerge from consecutive interactions within a global network, without the propagation of spikes along long-range axons; the features of such second-order traveling waves depend on local time constants within regions and conduction delays between regions. This framework naturally accommodates our observation of traveling waves propagating at modality-specific frequencies.

Due to the rotational nature of waves observed here, their anatomical pathways (starting and end points) cannot be determined from the present data. Auditory stimuli can reset the phase of ongoing oscillations in visual cortex^51,52^ and modulate subsequent visual responses^53^, but the routes through which the attention signal could propagate from the auditory system to visual cortex remain unclear. One possibility is a direct route. Direct anatomical projections from auditory cortex to visual cortex have been reported in non-human primates^54–56^. In humans, diffusion tractography suggested direct white matter connections between auditory and visual cortex^57,58^, complemented by resting-state functional connectivity between the two^59^. The other possibility is an indirect route relayed through the frontoparietal attention network^1^, which exerts top-down control over posterior alpha activity^60^. Several aspects of our data favor this second route. First, the iconic, binaural auditory cues require an arbitrary mapping from cue timbre to visuospatial locations that likely involves higher-order regions. Second, the lateralized traveling waves propagated from anterior to posterior over the dorsal scalp, a direction that matches a feedback signal from frontal regions to visual cortex. Third, the wave lateralization occurred several hundred milliseconds after cue onset, well after early cross-modal responses in visual cortex, which occur within about 50 ms^61^.

Together with causal^22,25^ and intracranial evidence^50^ for large-scale traveling waves, our findings suggest that attentional traveling waves are not an artifact or an epiphenomenon, but reflect inter-areal communication in the attention networks.

## STAR Methods

### Experimental model and study participant details

Twenty participants (14 females; mean age ± SD: 22.4 ± 4.0 years; range: 18-34 years) with normal or corrected-to-normal vision and no history of neurological or psychiatric disorders were recruited in the study. All participants were compatible with standard magnetoencephalography (MEG), EEG, and magnetic resonance imaging (MRI) procedures. Inclusion criteria were assessed by a medical doctor at a separate inclusion visit. The study was approved by the French Ethics Committee on Human Research (protocol number: 2017-A02787-46) and was performed at the Neuroimaging Center (CENIR) of the Paris Brain Institute (ICM) in full accordance with the Declaration of Helsinki (2013). All participants were naïve to the purpose of the study, provided written informed consent and were compensated for their participation. Two participants were excluded from the analysis due to failed attentional manipulation, resulting in a final sample of eighteen participants (12 females; mean age ± SD: 22.4 ± 4.3 years; range: 18-34 years).

### Method details

First note that in the present study, we only analyze the EEG and behavioral data. MRI and MEG data was also recorded for each participant and will be analyzed in a separate study.

#### Apparatus and Stimuli

Auditory stimuli were presented via MEG-compatible earphones, with a sampling rate of 44100 Hz. The experimental presentation was implemented using Psychtoolbox 3.0.18 in Matlab 2017b (Mathworks, Natick, Massachusetts)^62^ on a Windows 10 Professional computer. The spatially informative auditory cues were either a sine- or semisine-formed pure tone, while the neutral auditory cue was a sawtooth-formed pure tone (sampling rate of 44100 Hz, frequency of 1000 Hz, at comfortable volume). Each auditory cue was enveloped with 8-ms cosine rise and decay ramps. After training (see Training Session section), the informative cues were respectively associated with the spatial location (left or right) of the upcoming target grating stimulus. The specific associations between cues and locations were counterbalanced across participants.

Visual stimuli were displayed on a 79.5 x 44.5 cm screen (32.5 x 18.3 degree of visual angle, °VA) using a PROPixx projector (VPixx Technologies, Saint-Bruno, QC, Canada; 120 Hz refresh rate) positioned 78 cm away from the participant (resolution: 1920 x 1080 pixels). Two sinusoidal grating patches windowed by a raised-cosine (beta = 0.3) were presented, one in each lower quadrant of the screen (3°VA below the screen center, i.e., fixation, and 6°VA to the left or right of the screen center; 3°VA diameter, 2 cycles per degree, 20% contrast). One acted as distractor (vertical orientation) and the other as target (randomly tilted clockwise or counterclockwise from vertical; 25° maximum). The location of target and distractor gratings was counterbalanced across trials. Throughout the experiment, a white (RGB [255 255 255]) fixation cross (0.4°VA size, 0.1°VA linewidth) remained at the center of the screen.

#### Attention Cueing Task

Participants performed a two-alternative, forced-choice orientation discrimination task while covert, spatial endogenous attention was manipulated (**Figure 1A**). At the beginning of each trial, a 100ms-iconic auditory cue instructed participant to direct their attention towards the bottom right or left quadrant. After a variable delay (1350-1600 ms), two grating stimuli were simultaneously presented for 33 ms. In valid trials (66.6% of the trials), the cue was informative of the target location (tilted grating); attention remained sustained on the target. In invalid trials (33.3%), the target was at the opposite location from the cued one; attention had to reorient from the distractor to the target. After a 1500ms-delay, participants were instructed by a response cue (fixation turns black) to report the orientation of the target grating (clockwise or counterclockwise from vertical). Immediately after the button press or the maximum response window (1000 ms after response cue onset), feedback (fixation turns red RGB [255 0 0] for incorrect or no response, or green RGB [0 255 0] for correct) of 1000 ms was provided afterwards to keep participants motivated and focused. The inter-trial interval was jittered between 800-1300 ms.

The experiment had a within-participant design with two periods of interest: (1) post-auditory cue period in which sustained attention was directed by an auditory cue, and (2) post-visual stimulus period in which sustained attention was directed by a visual stimulus. Only valid trials were included in the analysis of post-visual stimulus period in the current study.

#### Experimental Procedures

Each participant underwent one training session, one concurrent MEG-EEG session and one session of anatomical and retinotopy MRI (for the purpose of another study; MEG, resting-state EEG and MRI data not used here). The MEG-EEG session consisted of MEG-EEG set up, breaks and experiment including staircase, main task, resting state and lasted a total of 4h. During this time, participants completed 10 5.5-min blocks (60 trials each) of the main task (400 trials valid and 200 invalid total) as described above. Participants were able to take short breaks between blocks and a longer break after 5 blocks.

##### Training session

A training session of 60 trials was performed during the week preceding the main experiment to help participants learn the association between the iconic cues and their corresponding spatial locations (left or right; counterbalanced across participants). Here, attentional cues were always valid, indicating the correct location of the upcoming target. The target grating was always tilted 25° clockwise or counterclockwise from vertical.

##### Staircase

At the beginning of the MEG-EEG session, a one-up-two-down staircase was used to tailor the specific target orientation to each participant to reach average performance of about 70% correct^63^. The initial orientation of the target grating was set at 20° from vertical, with a step size of 6°. Before the first reversal, a one-up-one-down rule was used. The step size was then divided by 2 every two reversals. The staircase block ended when the step size was smaller than 0.01°. The same staircase block was performed twice to familiarize participants with the task and ensure convergence of the threshold estimate (128 trials each maximum). The participant-specific discrimination target orientation was calculated by averaging the orientation from the last 20 trials in the second staircase block. To compensate for practice effects, the target orientation was then adjusted after each block of the main experiment to maintain accuracy at approximately 70% correct. The target orientation decreased from 5.72° ± 5.17° (mean ± SD across participants) after the staircase to 3.94° ± 2.57° in the last block.

#### Eye-tracking

Eye position was monitored throughout the main task to ensure central fixation (critical when manipulating covert attention) and avoid MEG-EEG signal artifacts. An infrared video camera (EyeLink 1000, SR Research, Ottawa, Canada) tracked the real-time position of the dominant eye. If a fixation break (blink, saccade or drift) of more than 10 ms within a 2.5°VA radius (does not overlap with any stimulus location) was detected between inter-trial interval onset and response cue onset, the trial would be aborted and repeated at the end of the block. This procedure maximized the number of analyzable trials.

#### EEG recording and preprocessing

EEG data was acquired simultaneously to the MEG using a 74-channel MEG compatible EasyCap, equipped with reference and ground electrodes, as well as electrodes for recording electrooculographic signals. Data collection was be conducted at a sampling rate of 1000 Hz, incorporating a high-pass filter at 0.01 Hz and a low-pass filter at 330 Hz to effectively isolate the frequency range of interest while minimizing external noise and signal artefacts.

EEG data were preprocessed and analyzed using custom-written scripts along with the MNE-Python toolbox^64^. Bad channels were detected manually by visual inspection and interpolated using MNE-Python defaults (spline interpolation). EEG recordings were offline re-referenced to the global average signal across all channels. EEG timeseries were first high-pass filtered at 0.1 Hz (causal FIR filter, Hamming window, minimum phase)^65^, and then low-passed filtered at 160 Hz (causal FIR filter, Hamming window, minimum phase) to avoid aliasing during subsequent decimation. Pre-processed EEG data were epoched from -2 s to 4 s around cue onset and then decimated from 1000 Hz to 500 Hz. Epochs were excluded if their amplitude exceeded a threshold of 5e-4 V, or if no response was given, resulting in 581.7 ± 12.7 (M ± SD) trials per participant (388.4 ± 8.1 valid trials).

#### Alpha power lateralization

Time-frequency decomposition was performed on single-trial data using Morlet wavelet convolution (two-cycle, 18 logarithmically spaced center frequencies between 2 Hz and 32 Hz). We choose this method, rather than for example FFT, because the events of interest are short-lasting, and can be washed out by frequency analysis over longer time windows^66^. For each trial, frequency, and time point, power was averaged across electrodes of the left and right hemispheres separately (32 electrodes per hemisphere; midline electrodes excluded). The Attention Modulation Index (AMI) was computed as:

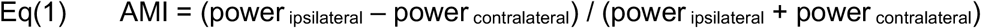

A positive AMI indicates higher power over the hemisphere ipsilateral to the attended location, that is, cued location after auditory onset and target location after visual onset.

To visualize the spatial distribution of alpha lateralization, alpha power was averaged over the significant clusters of AMI (0.690-1.020 s after auditory onset; 0.498-0.784 s after visual onset), separating left and right attended trials:

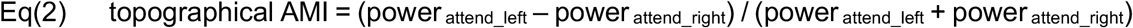

For each pair of electrodes, topographical AMI in the right hemisphere was subtracted from the left hemisphere, yielding a single hemispheric lateralization value displayed in the left hemisphere (see **Figure 1C and 1D**, right panels). Topographies are shown for descriptive purposes.

#### Traveling wave analysis

We identified traveling waves using a data-driven approach of dimensionality reduction (SVD)^25,67^. Traveling waves were analyzed from the complex wavelet coefficients (see Alpha power lateralization) using the WaveSpace toolbox^68^. Coefficients were first normalized to unit magnitude, so that all subsequent analyses were based on complex-valued phase alone. To identify the dominant spatial patterns of phase dynamics in our data, we applied SVD to decompose the complex phases across electrodes (pooled over frequencies, trials, and time points) into spatial components ordered by the proportion of variance they explain. The dominant components resembled a one-cycle rotating wave in the sagittal plane (**Figure S1**), consistent with previous reports of attention-related traveling waves propagating predominantly from anterior to posterior on the dorsal scalp^21,22,25^.

To facilitate interpretation and comparison across participants and conditions^25^, we then constructed rotational templates: each electrode was projected onto the sagittal plane, and its phase was given by its angular position around a singularity, such that phase propagates uniformly for one full cycle around the singularity. The singularity was located on the coronal plane through the preauricular points (y = 0), 10 cm (post-auditory cue period) or 5 cm (post-visual target period) above the interaural line, that is, inside the head beneath the vertex; these locations were chosen to best match the dominant SVD components in each period. Phase thus rotated around an axis parallel to the interaural axis. The clockwise (CW) template was defined as viewed from the left, and the counterclockwise (CCW), also viewed from the left, was its complex conjugate. Over the dorsal scalp, CW and CCW rotations corresponded to anterior-to-posterior and posterior-to-anterior propagations, respectively.

For each frequency, trial, and time point, the unit-magnitude complex-valued phases were projected onto each wave template after subtracting the mean of each electrode across frequencies, trials, and time points, yielding a complex coefficient β. A model phase pattern was obtained by rotating the template by the phase of β, and wave activity in each direction was quantified as:

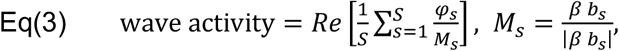

where φ_s_ is the phase at electrode s (before mean subtraction), and b_s_ is the template value at electrode s, | | denotes the magnitude of a complex number, applied separately at each electrode, and S is the number of electrodes. Because both φ_s_ and M_s_ have unit magnitude, this wave activity equals the mean cosine of the phase difference between the observed and modeled patterns across electrodes. Wave activity ranges from 0 to 1, with 1 indicating that the spatial phase pattern exactly matches the wave template and can be considered as a surrogate for spatiotemporal power in that direction.

#### Traveling wave lateralization

To test whether traveling waves were lateralized like alpha power, wave activity had to be estimated separately in the hemispheres ipsilateral and contralateral to the attended location. We therefore restricted the whole-brain templates to the electrodes of each hemisphere (midline electrodes excluded) and fitted them separately to the data of that hemisphere, using the same procedure as above. Because the templates rotate around the interaural axis, each hemisphere’s template retained a full rotation cycle, and the left and right templates were identical in shape. Hemispheres were labeled ipsilateral or contralateral relative to the attended location, and a Wave Modulation Index (WMI) was computed for each trial, frequency, and time point as:

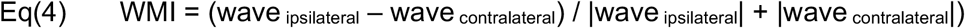

where wave_ipsilateral_ and wave_contralateral_ denote wave activity in the ipsilateral and contralateral hemispheres. The WMI was computed separately for CW and CCW waves.

#### Single-trial correlation analysis

To test whether traveling wave lateralization correlated with alpha power lateralization, we computed within-participant correlation coefficients across trials. For each trial, the WMI of CW waves was averaged within the theta band (auditory) or the alpha band (visual), and the AMI was averaged within the alpha band only. Both measures were computed from the same trials and averaged over a common time window, the union of significant clusters found above (auditory: 0.548-1.020 s; visual: 0.498-0.784 s). For each participant, the Pearson’s correlation between the two measures was computed across trials.

### Quantification and Statistical Analysis

#### Behavioral Analysis

We compared the behavioral performance in valid and invalid trials with two dependent variables, i.e., d’ and reaction time. D’ is a measure of sensitivity taking bias into consideration^69^. Individual and averaged d’ was calculated as:

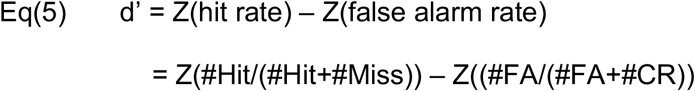

Higher sensitivity for valid than invalid trials confirmed successful attentional manipulation^70,71^. Shorter or equal median reaction time in valid than invalid trials excluded the possibility of speed-accuracy trade-off^26,27^.

#### Cluster-based permutation tests

Time-resolved lateralization indices, i.e., AMIs and WMIs, were tested against zero with one-sample cluster-based permutation tests (5000 permutations, one-tailed)^72^ over -0.1-1.5 s relative to auditory or visual onset.

#### Repeated-measures ANOVA

To characterize the hemispheric asymmetry of traveling waves, a 2×2 repeated-measures ANOVA with factors hemisphere (ipsilateral vs. contralateral) and direction (CW vs. CCW) was performed separately on theta-band wave activity after auditory onset and alpha-band wave activity after visual onset.

#### Correlation statistics

The within-participant Pearson’s correlation coefficients were Fisher z-transformed and tested against zero with a two-sided one-sample t-test.

#### Resource availability

All data and analysis code will be made publicly available upon acceptance of the manuscript. The MEG_Chicken toolbox (https://github.com/DugueLab/MEG_Chicken) was used by the first author as a training program for artifact detection (combined with training from the senior authors). The WaveSpace toolbox (https://github.com/kpetras/WaveSpace)^68^ was used for time-frequency decomposition and traveling wave analysis.

## Acknowledgements

This project has received funding from the European Research Council (ERC) under the European Union’s Horizon 2020 research and innovation programme (grant agreement No 852139 - Laura Dugué).

## Author contributions

L.D., K.P. and Y.K. designed the research. Y.K. performed the analysis of the data. Y.K., D.M.A and L.D. interpreted the data. Y.K. wrote the first version of the manuscript. All authors revised and edited subsequent versions of the manuscript.

## Supplementary Materials

**Figure S1.**
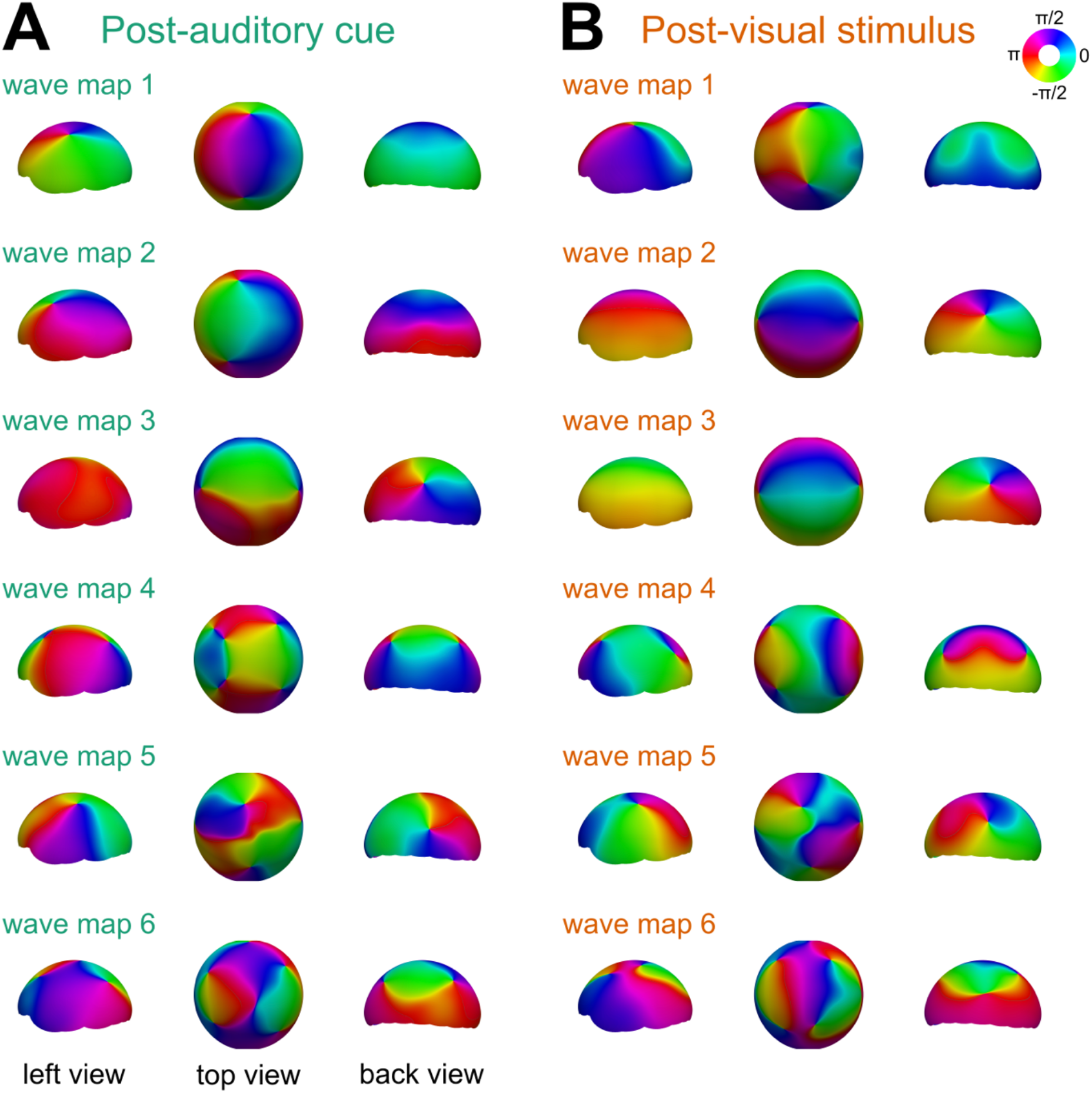
Dominant components of phase patterns extracted from the data. **A** The first six spatial basis vectors (wave maps) extracted in the post-auditory cue period, shown in the left, top and back views, respectively. Color indicates oscillatory phase. **B** same as **A**, in the post-visual stimulus period.

